# Lipid Disbalance Affects Neuronal Dendrite Growth and Maintenance in a Human Ceramide Synthase Disease Model

**DOI:** 10.1101/2024.10.31.621235

**Authors:** Anna B Ziegler, Cedrik Wesselmann, Konstantin Beckschäfer, Anna-Lena Wulf, Neena Dhiman, Peter Soba, Christoph Thiele, Reinhard Bauer, Gaia Tavosanis

## Abstract

The brain is susceptible to disturbances in lipid metabolism. Among the rare, genetically-linked epilepsies Progressive Myoclonic Epilepsy Type 8 (PME8), associated with the loss of Ceramide Synthase (CerS) activity, causes epileptic symptoms accompanied by neurodegenerative traits. We show that expression of a disease-causing c*erS* allele in *Drosophila* sensory neurons yielded developmental and degenerative dendrite loss. In *cerS* mutants, C18-C24 ceramides and membrane-forming complex sphingolipids, into which ceramides are converted, were reduced. At the same time bioactive signaling lipids including (dh)Sphingosine-1-P, deriving from the CerS substrate, were increased. To clarifying the etiology of PME8, we thus performed *in vivo* experiments to cell-autonomously rescue the individual metabolic alterations. We report that restoring specific long-chain ceramides while in parallel decreasing (dh)Sphingosine-1-P fully rescues the *cerS* mutant phenotype. Thus, despite the complex metabolic alterations, our data provide essential information about the metabolic origin of PME8 and delineate a potential therapy.

## Introduction

60% of the total dry mass of the human brain consists of lipids (1). Lipids are involved in a variety of biological processes. They are used as energy substrate, as building blocks for membranes, or function as bioactive signaling molecules (2). Because of the various and sensitive roles of individual lipids, neurons are highly vulnerable to alterations in their homeostasis. Ceramide metabolism deserves specific attention in this context, since a failure of ceramide synthesis or degradation results in a set of severe diseases with low prevalence, such as Progressive Myoclonic Epilepsy Type 8 (PME8), Hereditary sensory neuropathy (HSN1), Niemann-Pick Disease, Fragile X-associated tremor/ataxia syndrome (FXTAS), or Gaucher Disease (3,4). Each of these diseases is associated with an imbalance in ceramide amounts leading primarily to neurological symptoms, which emphasizes the importance of a well-controlled ceramide metabolism in the nervous system (3,5). In terms of their structure, ceramides are the simplest members of the large sphingolipid (SL) family. Ceramide Synthases (CerSs) are at the center of ceramide synthesis and thus also of complex SL production (6). Loss of CerS activity leads to PME8, a disease with an early onset (between 1 and 16 years of age) in which patients develop myoclonic seizures that appear to be related to loss of CerS activity in glia (7,8). In addition, PME8 patients suffer from neurodegeneration, as they develop moderate to severe and progressive cognitive impairment, ataxia, severe gait disturbances, and can become wheelchair bound (9,10). Interestingly, problems in the *de novo* ceramide synthesis pathway are frequently also associated with polyneuropathic phenotypes such as paresthesia, shooting pain, or disturbed pain delusions in the extremities (11,12). Whole exon sequencing and homozygosity mapping in patients identified two point mutations within *cerS1* (*cerS1*^H183Q^ and *cerS1*^R255C^) and a deletion on chromosome 1q21, which removes the entire *cerS2* gene (9,10,13,14). However, to date, the consequences of CerS loss on cellular lipid metabolism have not been adequately explored, and there is neither a cure nor an effective treatment for PME8 (15–18).

Ceramide *de novo* synthesis generates a large fraction of cellular ceramides. The pathway entails four enzymatic reactions by which at first (dh)sphingosine ((dh)S) is produced and then acylated to (dh)ceramide through the sphingosine N-acyltransferase activity of CerS (5). The catalytic center of CerS lays within the lag1p motive and includes two adjacent histidines that are required for enzymatic activity; one of those is mutated in the PME causing human *cerS1*^H183Q^ allele (19,20). Mammalian genomes harbor six CerS encoding genes (*cerS1-6*), which differ in their expression pattern and substrate specificity, accepting only acyl-CoA species of a defined chain length (21). The region that defines the acyl-chain specificity maps to the loop between transmembrane domains (TMDs) six and seven harboring the second human mutation (*cerS1*^R255C^) (Fig. 1A) (20).

**Figure 1.**
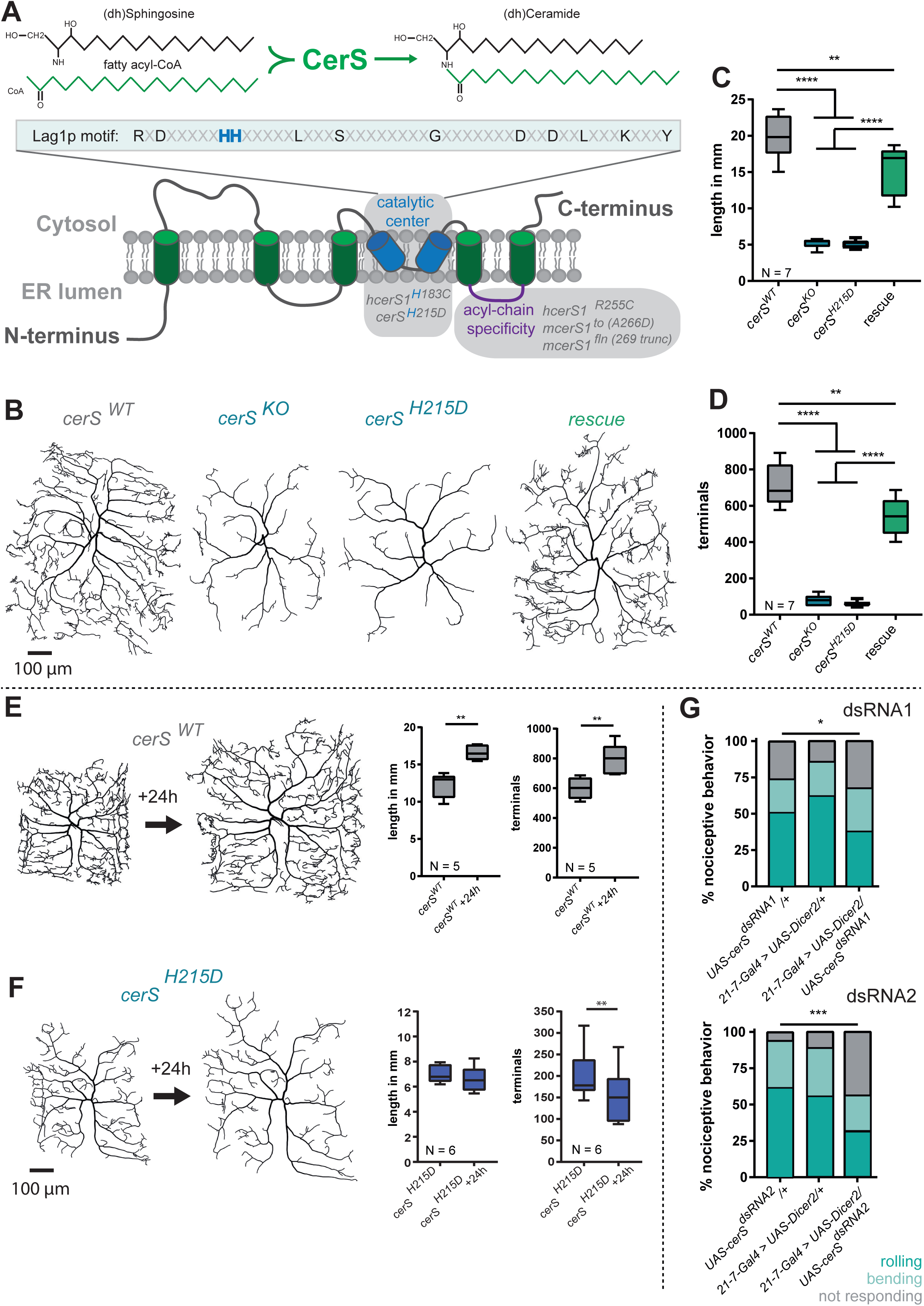
Cell-autonomous CerS activity is required for dendrite morphology: **A,** Up, CerS catalyzes the formation of (dh)ceramide from (dh)sphingosine and fatty Acyl-CoA. Down, schematic representation of CerS topology (modified after Thidar et al., 2018). The *hcerS1^H183C^* and the *Drosophila cerS^H215D^* mutations localize within the lag1p-motive that is required for CerS catalytic activity (blue). *hcerS1^R255C^* and m*cerS1^to^* mutations lay within a loop that determines acyl-chain specificity (purple). **B-D**, Mutant c4da single cell clones expressing wild-type CerS (*cerS^WT^*), no CerS (*cerS^KO^*) or the catalytic dead *cerS^H215D^* were generated using the MARCM technique. *CerS* expression was cell-autonomously rescued in *cerS^KO^* mutant neurons using a *UAS-cerS* expression construct (*cerS^KO^, UAS-cerS*). **B** Representative tracings of the single-cell MARCM clones. **C** and **D** Total dendrite length and total number of dendritic terminal endpoints, respectively. **E and F**, *CerS^WT^* and *cerS^H215D^* c4da MARCM clones were imaged in early LIIIf stage larvae. Afterwards, larvae were allowed to develop for 24h and the same neuron was re-imaged. Images show representative dendritic trees and graph the quantified total dendrite length and total number of dendritic terminal endpoints, respectively. **G,** Nociceptive responses (rolling and bending) of animals, in which *cerS* was knocked down using RNAi were assayed using a mechanonociceptive assay with a 50 mN von Frey filament. Statistical significance is indicated as follows in this and all following figures: *= p ≤ 0.05; **= p ≤ 0.01; ***= p ≤ 0.001; ****= p ≤ 0.0001. Statistical test used in C and D: 1-Way-ANOVA followed by Dunnett’s multiple comparison test; Statistical test used in E and F: paired student’s *t*-test.

Ceramides can be directly incorporated into membranes, or they can be used as substrate to produce complex SL. While the most prominent SLs in vertebrates are sphingomyelins (SM), invertebrates have high levels of ceramide-phosphatidylethanolamine (Cer-PE) (22). Ceramides with a specific acyl-chain length can also directly interact with ceramide-binding proteins and thereby fulfill various roles as bioactive compounds (23,24). Importantly, not only ceramides themselves but also the substrate for the CerS reaction, (dh)S, and its related metabolites (dh)S1P, Sphingosine (S) and Sphingosine-1-P (S1P), are lipid messenger molecules. Their action is largely linked to growth regulation and control of survival versus apoptosis (25). In the mouse brain, all of these so-called long-chain bases (LCBs) accumulate upon CerS dysfunction (18). Two mouse mutant lines (*toppler* (to) and *flincher* (fln)) harbor mutations in the acyl-chain binding loop of CerS1, resulting in viable but small adults suffering from ataxia (17,18). These studies established a link between general LCB accumulation and dendritic degeneration of Purkinje cells. However, also the lack of complex SL might also contribute to neurodegeneration in *cerS* mutant neurons (17,18). Typical for lipid metabolism-related diseases, it therefore remains unclear which of the complex alterations in lipid balance associated with loss of CerS function promotes neurite loss and degeneration and how PME8 could be potentially treated.

In contrast to mammals, the genome of invertebrates such as *Drosophila melanogaster* encodes for only one CerS, which is proposed to have broad substrate specificity (26). Reduction of CerS function enhances Fragile X-associated tremor/ataxia syndrome-linked neuronal degeneration in the fly eye (4). Furthermore, knock-down of *cerS* in *Drosophila* ventral lateral neurons (LNv’s) reduces their dendritic mass (27), suggesting a conserved role of CerS in neuronal dendrite complexity. These studies, together with the structural and functional conservation of this enzyme, suggest that *Drosophila* could be a useful model to understand the origin of the dendrite loss and neurodegeneration associated with *cerS* mutations. However, to date specific metabolic changes in fly *cerS* mutant neurons and their individual impact on neuronal morphology are unknown.

Here, we report loss of dendrite complexity in a newly generated *Drosophila* allele that mimics the PME8 causing human *cerS1*^H183Q^ mutation. We show that the abundance of multiple lipid species was changed in *Drosophila cerS* mutants. With quantitative genetic analysis, we furthermore demonstrate the impact of the individual lipid alterations on neuronal structural complexity. Finally, we identify the loss of long-chain ceramides and the accumulation of specific intermediates as key factors to maintain neuronal integrity and thus delineate paths for potential treatment.

## Results

### Cell-autonomous CerS activity is required for dendrite maintenance

To elucidate the impact of CerS dysfunction in *Drosophila* neurons, we first generated a new *cerS* allele that mimics the PME8 causing human *cerS1*^H183Q^ mutation. To do so we used a recently generated *cerS^KO^* allele in which most of the protein coding exons were replaced by an attP landing site (28). In a second step, this landing site was used to re-insert the missing coding exons of wild type *cerS* (*cerS^WT^*) (28) or the *cerS^H215D^* allele using ΦC31 integrase. The obtained *cerS^H215D^* allele contains the restored *cerS* coding sequence in its native locus, but carries a mutation in the lag1p motive that mimics the human disease-causing *cerS1^H183Q^* allele (Fig. 1A) (9,19,28). *cerS^KO^* and *cerS^H215D^* animals as well as the previously characterized strong hypomorphic allele *cerS^G0349^* died during early larval stages, which morphologically correspond to the first larval instar (26). Since the *cerS^H215D^* mutation caused early larval lethality, we could not directly test its impact on CerS enzymatic activity. However, we have previously shown that early developmental lethality of *cerS^G0349^* could be restored by expression of *UAS-cerS* but not upon expression of UAS-*cerS^H215D^* in the fat body. Also, overexpression of *UAS-cerS^H215D^* in wild-type larvae did not increase ceramide levels while overexpression of *UAS-cerS* did (26), supporting the idea that *cerS^H215D^* is enzymatically inactive.

In mice and flies loss of CerS function leads to reduction of neuronal dendrite complexity (18,27). We thus sought to characterize quantitatively the impact of the newly generated *cerS^H215D^* allele on dendrite complexity using the multimodal nociceptive class four dendritic arborization (c4da) sensory neurons of the fly larva as a cellular model (29). To overcome lethality we used the MARCM (Mosaic Analysis with a Repressible Cell Marker) technique to generate *cerS^WT^*, homozygous *cerS^KO^* or homozygous *cerS^H215D^* mutant c4da neuronal cell clones in an otherwise heterozygous background (30,31). In comparison to *cerS^WT^* c4da neurons, *cerS^KO^* or *cerS^H215D^* mutant neurons lost approximately 75% of total dendritic length and almost 90% of their total dendritic terminal endpoints (Fig. 1B-D). To test if only complex neuronal dendrites were affected by loss of CerS function, we analyzed the simpler dendrites of c1da and c3da *cerS* mutant neurons. Similarly, c1da and c3da neurons displayed a clear reduction in the number of dendrites (Suppl. Fig 1A/B), suggesting that CerS function may in general be necessary for neurons to properly build or maintain their dendrites. We next confirmed that the observed phenotype was due to the specific, cell-autonomous loss of CerS activity by re-expressing *UAS-cerS* (32) in the *cerS^KO^* mutant da neurons. This rescued total dendrite length and the total number dendrite terminal endpoints to a large extent (Fig. 1B-D and Suppl. Fig. 1A/B).

To distinguish between neurodevelopmental growth defects and neurite loss in the *cerS* mutants, we compared the dendrites of mutants and controls at several developmental stages. Mutant c4da neurons showed already significant dendrite growth deficits at the second instar larval stage which strongly suggests an early developmental component (Suppl. Fig. 1C). In addition, though, the difference in dendritic tree morphology between mutants and control became more pronounced during the third larval instar (LIII), which is divided into a feeding period (LIIIf) and the later wandering stage (LIIIw) (Suppl. Fig. 1C). *cerS^WT^* c4da dendrites constantly grew in length and gained in dendritic complexity, whereas *cerS^KO^* mutant c4da neurons showed an even simpler dendritic tree morphology at the LIIIw stage than in the younger LIIIf stage. These data suggest that here, in addition to the early developmental defects, degenerative processes might be at play. To specifically address the progressive loss of dendrite branchlets during the LIII stage we have imaged *cerS* deficient MARCM clones at the early LIIIf stage and re-imaged the same cells after 24h. This analysis confirmed the progressive loss of identifiable dendrites in *cerS^H215D^* mutant c4da neurons (Fig. 1E/F) or in *cerS^KO^* mutant c4da neurons (Suppl. Fig. 1D). Loss of *cerS* was suggested to induce caspase activity in cultured mouse hippocampal neurons (18). We therefore tested whether caspase activity was also involved in the loss of dendrites of *cerS* mutant c4da neurons. However, no caspase activity could be detected using Apoliner, a caspase activity reporter (33) (Suppl. Fig. 2A). Furthermore, the expression of the caspase inhibitor p35 in mutant c4da neurons did not rescue the dendrite defects in *cerS* mutant c4da neurons (34) (Suppl. Fig. 2B/C). Lastly, we could follow individual *cerS^KO^* c4da mutant MARCM clones from the third larval instar to the late pupal stage (Suppl. Fig. 2D/E), indicating that the neurons survive during this period. Taken together, differently to what has been observed in cultured neurons (17), *in vivo* we described severe morphological alterations, which, at the observed stages, are not related to caspase-induced apoptosis. To finally explore whether the altered c4da dendrite morphology was associated with functional response defects, we performed a mechano-nociceptive assay to measure the c4da neurons response to noxious stimuli (35,36). RNAi-mediated knockdown of *cerS* expression in c4da neurons resulted in significantly less nociceptive responses compared to controls using two different *UAS-cerS^dsRNA^* lines (Fig. 1G) suggesting *cerS* is also required to maintain c4da neuron function.

### Impairment of the Cer-PE *de novo* synthesis pathway leads to dendrite simplification in c4da neurons

To address the specific changes in lipid composition caused by lack of *cerS* function, we analyzed the lipidome of *cerS* mutant animals using shotgun tandem mass spectrometry (MS). CerSs are most prominently involved in ceramide *de novo* synthesis and hence in the production of complex SL such as Cer-PE (Fig. 2A). *cerS^KO^* or *cerS^H215D^* homozygous mutant animals die very early in development as small larvae, which morphologically correspond in size to first instar. We thus used the weaker *cerS^P61^* allele, in which the *cerS* mRNA level was reduced by approximately 60% and which gave rise to a low number of slim larvae reaching the LIII stage (26). As previously reported (26), we found a reduction of the total lipid level and consequently also of the absolute ceramide levels (Fig. 2B left, and Suppl. Fig. 3A). We next measured the relative ceramide levels within the remaining lipid fraction and found them not altered in the mutant. Since ceramides are used to produce complex sphingolipids, we also measured the relative levels of Cer-PE, which is the major complex sphingolipid in *Drosophila*, and found it to be decreased (Fig. 2B, right). We next examined how a defective Cer-PE production would affect the morphology of c4da neurons and analyzed the dendritic morphology of single cell clones generated by MARCM bearing mutations that blocked the Cer-PE producing pathway at different stages. To this end, we compared the dendrite morphology of c4da neurons carrying loss of function mutations for *lace* (*lace^SK6^*), *des1* (*des1^KO^*), or *cpeS* (*cpeS^KO^*) with the effect of the loss-of-function allele *cerS^H215D^*. We exploited the highly quantitative morphological properties of c4da neurons to be able to compare the impact of each of these mutations and found that neurons carrying mutations at each of the steps of the Cer-PE producing pathway display reduced dendritic complexity (Fig. 2A, C). However, loss of *cerS* activity in c4da neurons resulted in the quantitatively strongest reduction (Fig. 2D-E). We verified this result by knocking down the Spt1 complex members *spt1* and *lace* as well as *3-ksr, cerS, des1,* or *cpeS* in c4da neurons using RNAi. Similarly to the outcome of the MARCM approach, the dendrite morphology of c4da neurons was most strongly affected by knocking down *cerS* (Suppl. Fig. 3B-D). Taken together, these data indicate that Cer-PE production is essential to establish complex dendrite morphologies. In addition, they suggest that multiple intermediate species can be important for c4da neuron dendrites. However, the fact that impairment of the function of enzymes up- or downstream of *cerS* leads to milder phenotypes than impairment of *cerS* function itself suggests that the strong phenotype in *cerS* mutant neurons may be due to additional metabolic problems.

**Figure 2.**
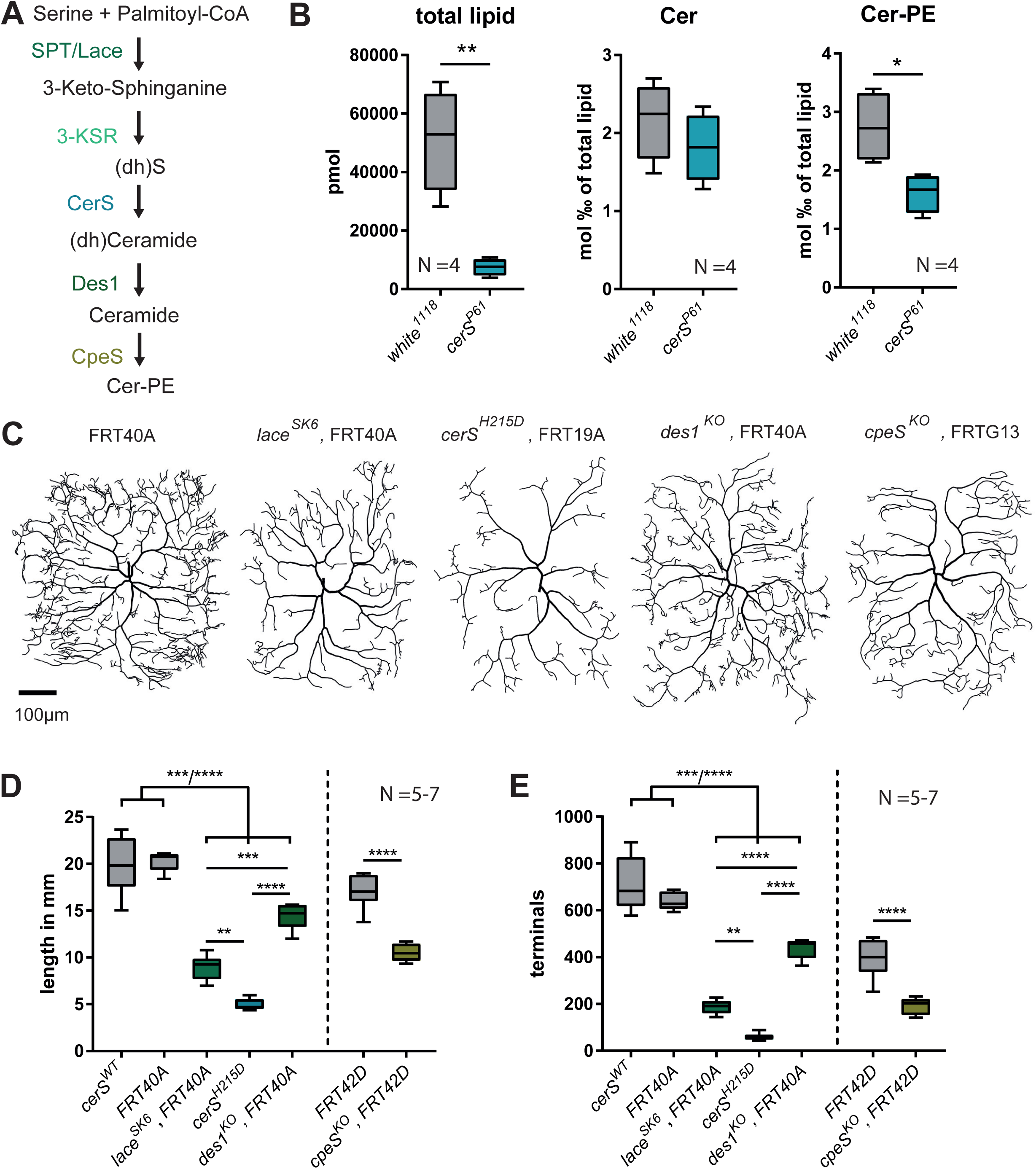
*cerS* deficiency strongly affects dendrite morphology: **A,** Schematic representation of the Cer-PE synthesis pathway. **B,** Lipids were extracted from control larvae (*white^1118^*) or *cerS^P61^* mutant escapers at the LIII stage and lipid levels were measured by tandem shotgun MS. Graphs show total lipid levels (left) and relative ceramide (middle) and Cer-PE (right) levels. **C-E**, Single homozygous *lace^SK6^*, *cerS^H215D^*, *des1^KO^* or *cpeS^KO^* mutant c4da neurons were generated using the MARCM technique. Quantified total dendrite length and total number of dendritic terminal endpoints are shown in **D** and **E**, respectively. Representative neuronal tracings are shown in **C.** Statistical test used in B: paired student’s *t*-test, D and E: 1-Way-ANOVA followed by Dunnett’s multiple comparison test.

### Elevated (dh)S1P levels negatively affect dendrite morphology

The Spt-complex generates (dh)S which is the substrate for the CerS reaction. Given the milder phenotype of the Spt-complex member *lace* in comparison to the *cerS^H215D^* mutation, we tested if the accumulation of (dh)S or its phosphorylated metabolite (dh)S1P, could also contribute to the impact of *cerS* mutation on neuronal morphology (Fig. 3A). Indeed, elevation of LCBs like Sphingosine, (dh)S, S1P, and (dh)S1P was previously already correlated with a decreased dendrite complexity (16–18). LCBs are bioactive signaling molecules and are involved in signaling pathways regulating survival, apoptosis and proliferation (25,37,38). To address the impact of LCB accumulation on dendrite morphology of c4da neurons, we first asked which of these LCBs accumulated in *cerS* mutant fruit flies. As above, we used hypomorph *cerS^P61^* mutant larvae for these experiments. In these animals, we measured increased levels of (dh)S and (dh)S1P (Fig. 3B). Since the accumulation of the phosphorylated and/or of the non-phosphorylated metabolite influences neuronal viability (16,17) we tested which of the two species has greater influence on c4da dendrite morphology. To do so, we generated *cerS^H215D^* mutant neurons in MARCM experiments in which we simultaneously expressed sphingosine kinases (SK) so that accumulated (dh)S will be preferentially converted to (dh)S1P (Fig. 3A). The *Drosophila* genome encodes for two functionally redundant SKs (SK1 and SK2); their overexpression enhances the production of (dh)S1P from (dh)S and induces neurodegeneration of photoreceptors in adult *Drosophila* (16,39). In c4da neurons expression of either of the two SKs strongly enhanced the loss of dendrites in *cerS ^H215D^* deficient cells (Fig. 3C-E). By contrast, expression of Sphingosine-1-phosphate lyase (Sply), which degrades (dh)S1P to a fatty aldehyde and phosphoethanolamine, mildly rescued the dendritic phenotype (Fig. 3C-E), confirming that enhanced (dh)S1P levels are detrimental for neuronal dendrite morphology. Taken together, while phosphorylated LCBs have been frequently described as anti-apoptotic factors (40), our data suggest that an increased level of phosphorylated LCBs in neurons leads to a strong morphological simplification and involve these species in the detrimental impact of loss of CerS function.

**Figure 3.**
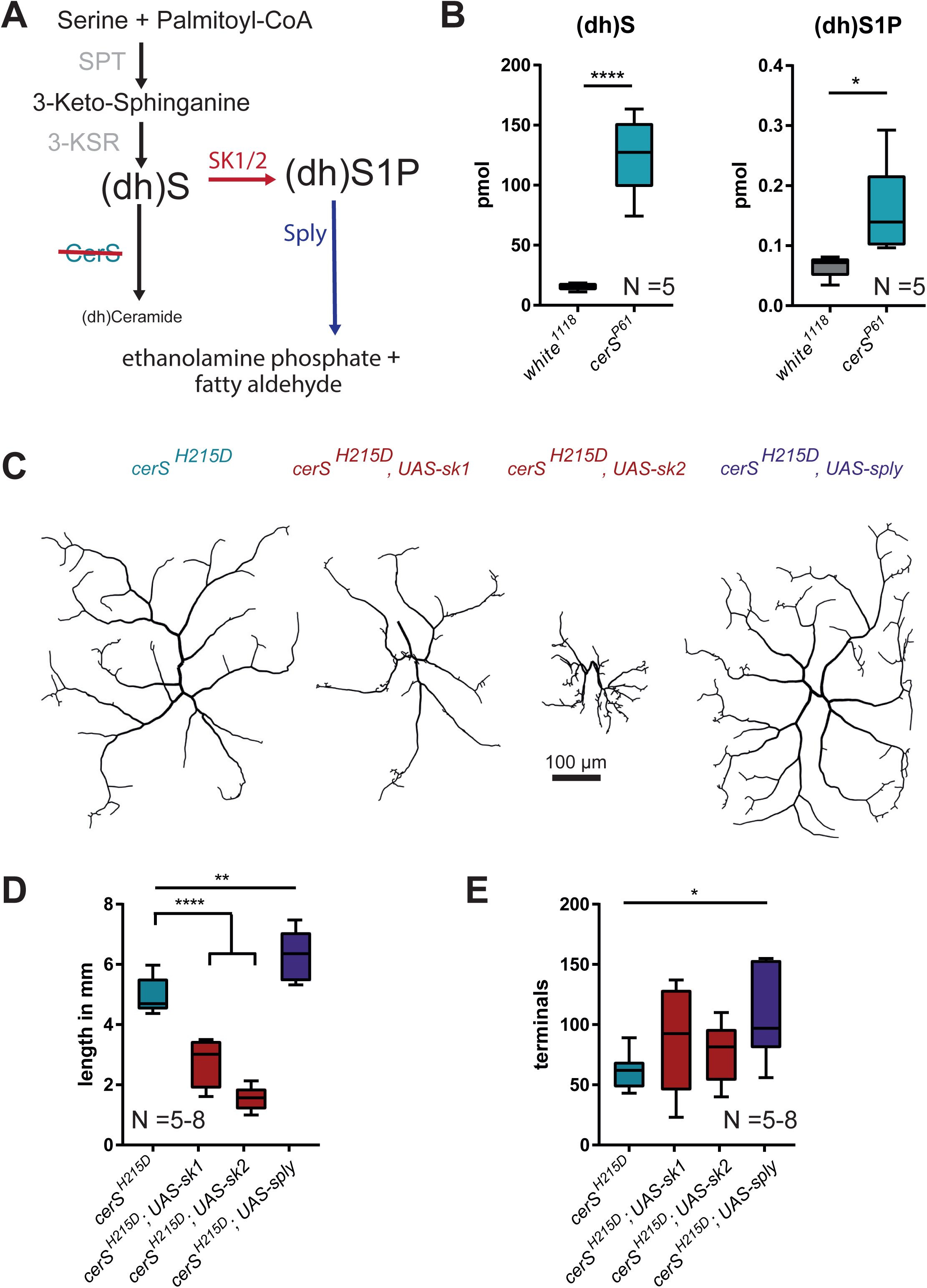
Elevated (dh)S1P levels negatively alter dendrite morphology: **A,** (dh)S accumulates in *cerS* mutant cells and is converted to (dh)S1P by SK1 or SK2. (dh)S1P is degraded to ethanolamine phosphate and a fatty aldehyde by Sply. **B,** absolute levels of (dh)S and (dh)S1P in control (*white^1118^*) and (*cerS^P61^*) mutant larvae. **C-E,** Single c4da MARCM clones homozygous *cerS^H215D^* mutant or homozygous *cerS^H215D^* mutant simultaneously overexpressing SK1, SK2, or Sply. Quantified total dendrite length and total number of dendritic terminal endpoints are shown in **D** and **E**, respectively. Representative neuronal tracings are shown in **C.** Statistical test used in B: paired student’s *t*-test and in D and E: 1-Way-ANOVA followed by Dunnett’s multiple comparison.

### C4da neurons dendrite complexity depends on C18-C24 ceramides supply

Out data so far indicated that both, the accumulation of (dh)S and (dh)S1P as well as the lack of ceramides for Cer-PE production are detrimental for c4da neuronal morphology. We thus reasoned that both metabolic alterations might need to be reverted to obtain a full rescue of the phenotype. As a next step towards designing efficient rescue strategies, we thus asked whether all ceramides were affected by the loss of *Drosophila* CerS function. We found that only ceramides with acyl-chains between C18-C24 carbons were reduced in the *cerS^P61^* mutant while surprisingly ceramides with shorter acyl chains (C12, C16) were not (Fig. 4A). This data indicate that *Drosophila* CerS produces ceramides with C18-C24 acyl-chains. They also suggest that ceramides with shorter acyl-chains are likely derived from alternative pathways or from an exogenous source such as food or gut bacteria. Interestingly, a shift in the ceramide species composition was also observed in *cerS1* deficient mice but the impact of an imbalance in ceramide species on the nervous system had not been investigated (17).

**Figure 4.**
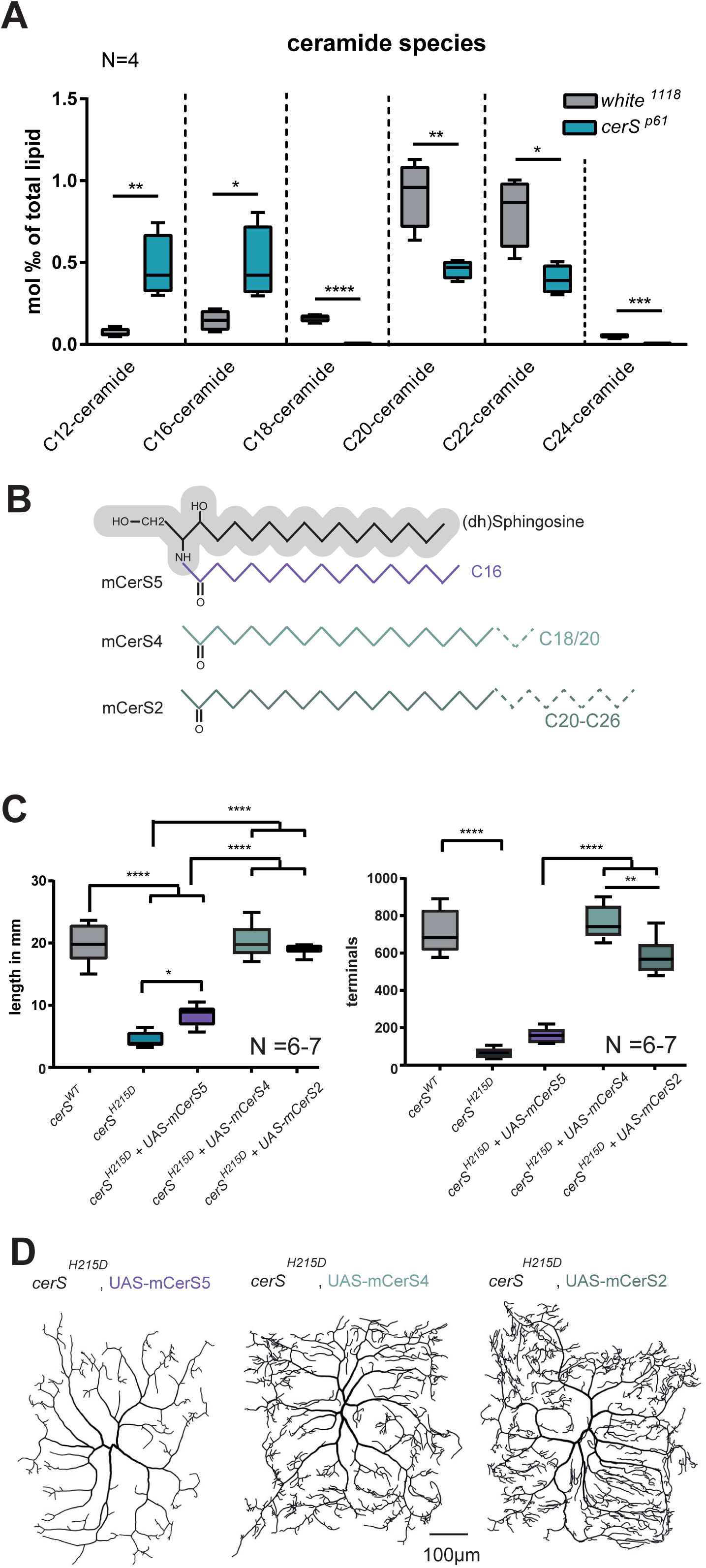
Neuronal morphology relies on a supply with C18-C24 ceramides: **A,** Lipids were extracted from LIII larvae and ceramide levels were measured by tandem shotgun MS. **B,** Schematic representation of acyl-chain substrate specificity of mCerSs. The (dh)S backbone is represented in gray and specific acyl-chains used by mCerS in color. **C-D,** mCerSs generating mostly C16-Cer (mCerS5), C18-C20-Cer (mCerS4), or C20-C26-Cer (mCerS2) were expressed to rescue the lack of endogenous CerS function in homozygous *cerS^H215D^* mutant c4da MARCM clones. Graphs show the quantified total dendrite length and total number of dendritic terminal endpoints, respectively **(C)**. Representative neuronal tracings are shown in **D.** Statistical test used in A: paired student’s *t*-test and in C: 1-Way-ANOVA followed by Dunnett’s multiple comparison.

The differential impact of the *cerS* mutation on the abundance of ceramides of different lengths prompted us to test the effect of restoring the levels of individual ceramide species in the *cerS* mutant background. To this aim, we took advantage of the specificity of mammalian CerSs towards acyl-chains of a given length to produce ceramides (Fig 4B). We thus created *cerS^H215D^* mutant c4da cell clones, in which we rescued the lack of the *Drosophila* CerS function expressing different murine CerS enzymes. By doing so (dh)S and (dh)S1P should be lowered and simultaneously only specific ceramides provided. The dendritic trees of mutant neurons expressing mCerS2 or mCerS4, which produce ceramides with longer acyl-chains (C18-C20 or C20-C28, respectively), were comparable to those of wildtype c4da neurons suggesting that these mouse enzymes can fully compensate for the loss of the *Drosophila* enzyme. By contrast, expression of mCerS5, which preferentially utilizes C16 fatty acyl-CoA, only minimally improves the dendrite complexity of *cerS* mutant c4da neurons. The complete rescue of the mutant dendritic phenotype by increasing ceramides with an acyl-chain of at least 18 carbons indicates that these ceramide species are key metabolites essential for the complex dendrite elaboration of c4da neurons (Fig 4C/D). To extend this finding and directly ask whether supplementation with long-chain ceramides might represent a path towards improving c4da neurons dendrite complexity, we fed larvae carrying single *cerS^H215D^* mutant c4da cell clones with C-16 or with C-18 ceramides. Also in this different context, supplementation with C-18, but not C-16, ceramides led to a mild but significant improvement of c4da dendrite complexity (Suppl. Fig 4).

Taken together, ceramides with acyl-chains longer than C16 promote c4da neurons’ dendrite complexity in a *cerS* mutant model. They furthermore represent a key supplement for restoring defects induced by *cerS* mutations.

## Discussion

Diseases that affect lipid metabolism and in particular, those directly linked to ceramide anabolic or catabolic pathways, frequently result in neurological diseases and neurodegeneration (3). Here, we demonstrated that CerS function is essential for the establishment and maintenance of dendrite complexity of c4da neurons. Combining genetic analysis and lipidomics, we showed that a dysfunctional CerS causes multiple alterations in lipid metabolism, such as the lack of membrane lipid production (Cer-PE), reduction of long-chain ceramides abundance and the increase of bioactive lipids such as (dh)S1P. Exploiting the quantitative character of c4da dendrite morphology, we defined the complex picture of lipid metabolism alterations associated with PME8-related mutations of CerS and delivered evidence that multiple of these individual changes have an impact on dendrite complexity in da neurons. Our data identify the accumulation of (dh)S1P and the loss of specific long-chain ceramides as key detrimental lipid species alterations. They furthermore point to supplementation of long-chain ceramides as a potential path towards restoring dendrite complexity in these mutants.

### Lack of complex sphingolipids affects c4da dendrite morphology

At first, we showed that the *cerS* mutation yielded a strong reduction of Cer-PE levels. Cer-PE is the most prominent invertebrate SL and the functional counterpart of mammalian sphingomyelin (SM). It is produced from ceramide by CpeS (7,22) and the strong reduction of the absolute ceramide levels in *cerS* hypomorphs can well explain its low levels in the mutants.

. In *Drosophila*, the lack of glial Cer-PE production in *cpeS* mutants was recently also shown to cause photosensitive epilepsy and perturbed circadian rhythm. *cpeS* deficiency during development resulted in animals that had morphologically defective glial cells and a rescue of *cpeS* expression in glia was able to rescue the above mentioned defects (7,41). In addition to providing membrane lipids, which serve as building blocks for the developmental cell membrane expansion (42), complex SLs in the plasma membrane, including Cer-PE and SM, along with glycosphingolipids and cholesterol, form lipid rafts with higher viscosity. Lipid rafts are required for the clustering of proteins or signaling molecules, for membrane trafficking, and interact with the submembrane actin cytoskeleton (43). A Cer-PE deficient nervous system fails to establish lipid rafts (7) which suggests that all above mentioned processes might well be altered in *cerS* deficient neurons, too, and thus could contribute to the aberrant c4da dendrite morphology.

### Dendrite morphology is strongly affected by elevated (dh)S1P levels

Altered LCB levels were frequently correlated to neuronal dysfunction. For example, a lack of LCB production in Spt-complex mutant *Drosophila* neurons was correlated with protein sorting defects and aggregation of cell recognition proteins, leading to axon morphology defects (44). Similar to what was published in *cerS* mutant mice (17) we found increased phosphorylated and non-phosphorylated LCB levels (Fig. 3B). A genotype-phenotype association study of Hereditary sensory and autonomic neuropathy type 1 (HSAN1) in patients also pointed towards a highly toxic effect of high LCB levels on the nervous system. HSAN1 is a polyneuropathy resulting from point mutations within the SPT complex, which functions upstream of CerS in the ceramide *de novo* synthesis pathway. Each of the 17 mutations mapped so far affect the peripheral nervous system but, compared to the symptoms of PME8 patients, with overall mild consequences, no seizures, and a later disease onset. Interestingly, though, three of the 17 mutations lead to increased enzyme activity and increase LCB levels- and these patients show exceptionally severe HSAN1 symptoms. Therefore, high plasma spingoid base levels were suggested as a biomarker that could help to predict the severity of the HSAN1 (11). However, which LCB metabolite is detrimental for neuronal viability could not yet been sufficiently explored. Here, we demonstrated that both (dh)S and (dh)S1P levels were increased in *cerS* mutants and revealed by genetic analysis that (dh)S1P seems to be the more toxic species (Fig.3). The effect of (dh)S1P had not yet been explored extensively in neurons, but its analog S1P acts on neural development, differentiation, migration, survival, and synaptic transmission (45,46). It is therefore not surprising that its concentration is tightly regulated by SKs and phosphatases like Sply. Phosphorylated LCBs are pleiotropic messenger molecules. They can for instance act trough S1P-receptors (47) and blocking this signaling pathway might represent a valuable approach to limit the loss of dendrites in *cerS* mutant c4da neurons. Unfortunately, fly S1P-receptors have not yet been identified. However, based on the data presented here, it will be of great interest to block (dh)S1P signaling using S1P-receptor inhibitors in mammalian PME8 models, in which S1P receptors were extensively characterized (40).

### Neuronal morphology relies on C18-C24 ceramides supply

When comparing the composition of ceramide species between the mutant and the control, we found that *cerS* mutants have an increased relative amount of C12- and C16-ceramides while C18-C24-ceramides were reduced, changing the relative abundance of each ceramide species. We thus propose that the fly CerS is responsible for the synthesis of longer acyl-chain ceramides (C18-C24), while C16-ceramides derive from an exogenous source such as the diet or gut bacteria (48,49). Interestingly, also in *cerS1* and *cerS2* mutant mice increased C16-ceramides levels were observed (17,18,23,50). Thus, imbalance in the relative abundance of ceramide species might be a conserved trait. However, it remained unclear whether the shift in ceramide species composition also contributed to the observed loss of Purkinje cell neurites in these mutants (18,51). Indeed, only recently attention has been paid to the effects of individual ceramide species, such as C16-ceramides, and their impact on cell vitality and survival (23,52). C16-ceramides can induce ER stress (24,52–54); in addition, in mitochondria C16-ceramides may form channels permeabilizing the mitochondrial outer membrane and inducing mitochondrial dysfunction (52,55,56).

We not only showed the differential impact of a *cerS* mutation on the abundance of different ceramide species. Our analysis of c4da neuron dendrites in the *cerS^H215D^* mutant background allowed us also to dissect the role of those individual ceramide species. We found that only CerS enzymes that synthetize long-chain ceramides (longer than C-16) can fully restore dendrite complexity and maintenance in these mutants. This finding supports the view that individual ceramide species display distinctive functions. It furthermore allows to hypothesize a path for improving the outcome of *cerS* mutations, including in PME8 patients. While our genetic analysis pointed to a cell autonomous requirement for ceramide synthesis in c4da neurons, we also found that mere dietary supplementation of the appropriate long-chain ceramides could improve the c4da neurons defects. We thus envision that a combination of reduction of toxic LCBs, such as (dh)S1P, or of their impact - together with supplementation of long-chain ceramides will be an important path to be tested in future experiments in vertebrate models of PME8.

In summary, we showed that the dendrites of *cerS* deficient neurons display reduced complexity already early in development and additionally progressively lose dendritic branches at later developmental stages. We dissected each of the multiple alterations in lipid metabolism that arise in *cerS* deficient cells and analyzed their individual quantitative impact on the morphology of c4da neurons. Overall, our analysis indicates the accumulation of LCBs, in particular (dh)S1P, and the reduction of long acyl-chain ceramide species (C18-C24-ceramides) as the key factors determining the neuronal phenotype- and points to paths for restoring neuronal integrity in this model of PME8 (Fig. 5).

**Figure 5.**
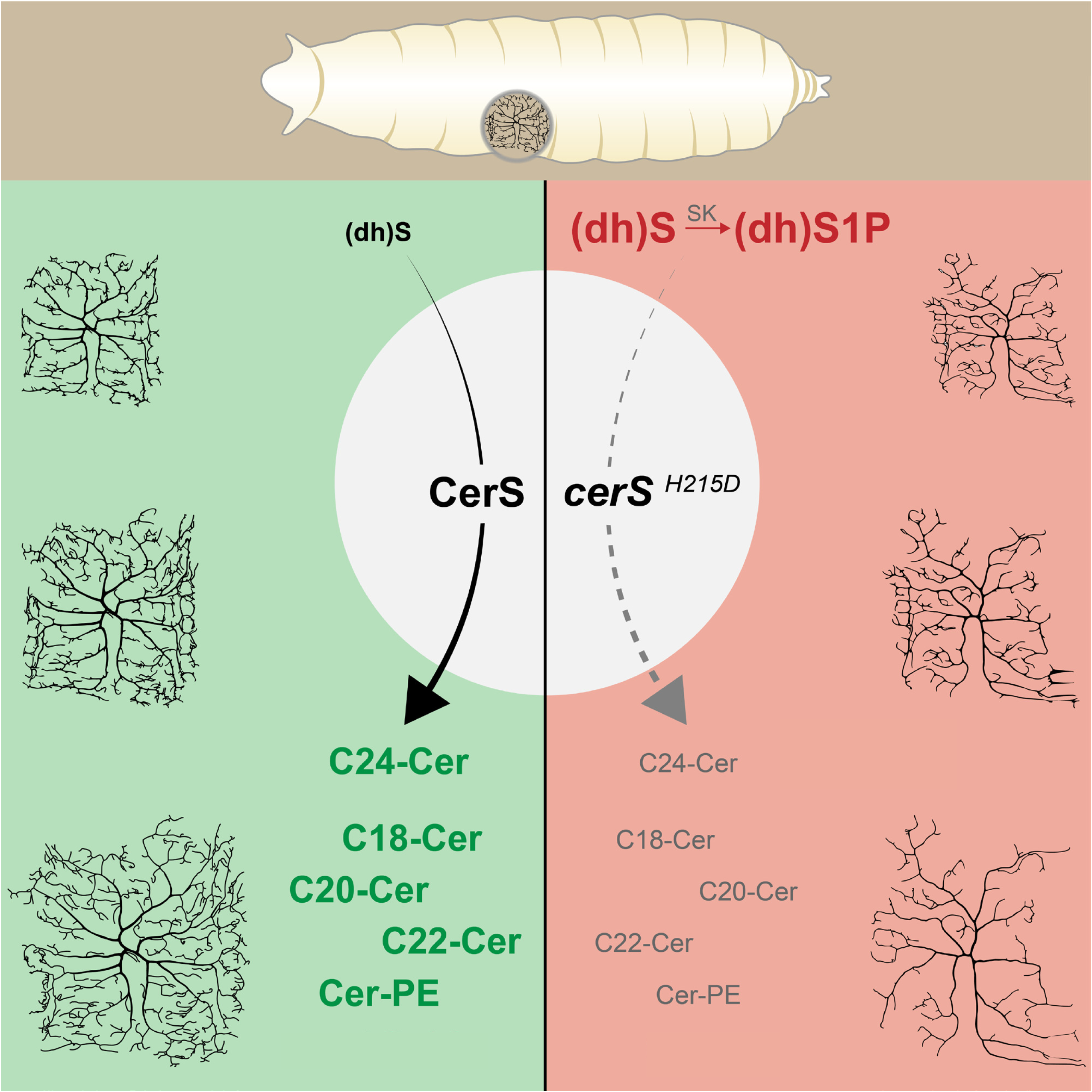
Model: CerS activity reduces LCBs such as (dh)S and (dh)S1P while producing long-chain ceramides (left). In *cerS* deficient neurons (dh)S and (dh)S1P levels are elevated and simultaneously C18-C24-ceramides diminished. To fully rescue neuronal vitality, both metabolic alterations must be reverted.

Furthermore, this work demonstrates the importance of a detailed lipidomic analysis combined with -genetic dissection of metabolic pathways, to formulate informed hypotheses on the origin and potential treatment of lipid-metabolism related diseases.

## Material and Methods

### Fly husbandry and maintenance

To generate MARCM clones, mutant *cerS* alleles *cerS^H215D^* (this work), *cerS^KO^* and the control allele *CerS^WT^* (28) were recombined with the FRT19A site. *lace^SK6^*, FRT40A was obtained from the NIG stock center (M2L-0156). This allele lacks 11 bp, which leads to a frame shift and resulting in a truncated protein. *des1^KO^*, FRT40A (57) was generously provided by Chin-Chiang Chan (National Taiwan University). MARCM clones were generated by using the SOP-FLP lines Nr. 109946 (FRT19A) and 109947 (FRT40A) (Kyoto stock center, Japan) (58).

For RNAi experiments we generated a fly line harboring *ppk-Gal4* (Chr. III) (kind gift from Yuh-Nung Jan, Howard Hughes Medical Institute, San Francisco, USA) recombined with *UAS-mCD8GFP* (Bloomington stock Nr. 5130) and *UAS-Dicer2* (Chr. II, kind gift from Justin Blau, New York University, USA) and crossed it to the following UAS-RNAi lines: *UAS-schlank^dsRNA1^* (Nr. 109418); *UAS-schlank^dsRNA2^* (Nr. 41115); *UAS-des1* (Nr. 106665); *UAS-lace^dsRNA1^* (21805); *UAS-lace^dsRNA2^* (110181); *UAS-spt1^dsRNA1^* (Nr. 10020); *UAS-spt1^dsRNA2^* (108833); *UAS-3KSR^dsRNA1^* (Nr.6734); *UAS-3KSR ^dsRNA2^* (109284), *UAS-OR42b^dsRNA^* (control, Nr. 101143) all obtained from the VDRC stock center in Vienna, Austria. Crosses were kept at 27 °C.

*UAS-schlank* (Nr. F000075) was obtained from the FlyORF collection Zürich Switzerland, *UAS-SK1* and *UAS-SK2* (16) were kindly provided by Usha Acharya, University of Massachusetts Medical School, US. *UAS-spin* was obtained from the Bloomington stock center, US (Nr. 39669) and *UAS-sply* (59) was generously provided by Corinne Antignac, Paris, France.

Flies were maintained on standard medium at 25°C, unless otherwise stated. Ceramides for the feeding assay were purchased from Avanti Research, US and used at a concentration of 0,16 mg/ml (C16-ceramide) and 0,17 mg/ml (C18-ceramdie).

### Generation of the *cerS^H215D^* mutant

*cerS* exons 2-6 were cloned into pGE-attB-*cerS^WT^* by Sociale et al 2018 (28,60). To generate pGE-attB-*cerS^H215D^* quick change PCR was used to replace histidine at position 216 by aspartate using pGE-attB-*cerS^WT^* and the following primers: cagatgttcatc**gat**cacatggtcacc and ggtgaccatgtg**atc**gatgaacatctg (bases encoding for histidine/aspartate are indicated in bold). *cerS^H215D^* transgenic flies *were* made as previously described. In brief, we used the previously published founder line (*cerS^KO^*) in which *cerS* exons 2-6 were removed by homologous recombination while simultaneously introducing an attP landing site (28,60). Next, the landing site was used to re-integrate the *cerS* encoding exons via ΦC31-mediated integration. The integration of pGE-attB-white^+^-*cerS^H215D^* into the white eyed *cerS^KO^* was done by BestGene Inc. (Chino Hills, CA, USA) and transgene integration was marked by White^+^ expression. The correct re-insertion of the *cerS* encoding sequence was tested by PCR. Since *cerS^H215D^* expression is derived from the endogenous locus it should follow the expression level of wildtype *cerS* which we confirmed via qRT-PCR.

### Live Imaging and Image Processing

The dendritic morphology of da neurons was observed by immobilizing living larvae between a glass slide and a coverslip in a mixture of halocarbon oil and ether (4:1). One da neuron per animal was imaged by confocal microscopy (Zeiss LSM700 or LSM710). Images were hand traced and analyzed using the TREES toolbox plug-in for MATLAB (R2014b) (61).

### Lipidomics

Lipid extraction: Lipids were extracted from individual larvae. Each larva was homogenized in 150µL water (LC-MS grade). To every homogenate, 1000µL Extraction mix (5/1 MeOH/CHCl_3_ containing internal standards. PE[31:1] 420pmol; PC[31:1] 792pmol; PS[31:1] 197pmol; PI[34:0] 169pmol; PA[31:1] 112pmol; PG[28:0] 103pmol; CL[56:0] 57pmol; LPA[17:0 79pmol; LPC[17:1] 70pmol; LPE[17:0] 76pmol; Cer[17:0] 64pmol; SM[17:0] 198pmol; GlcCer[12:0] 110pmol; GM3[18:0-D3] 29pmol; TG[50:1-D4] 2351pmol; CE[17:1] 223pmol; DG[31:1] 128pmol; MG[17:1] 207pmol; Chol[D6] 1448pmol; Car[15:0] 91pmol; Sph[d18:0-D7] 64pmol) were added and tubes were sonicated for 30 minutes in a bath sonicator. After 2 minutes centrifugation at 20000g, supernatant was transferred to a fresh Eppendorf tube. 200 µL chloroform as well as 600µL 1% acetic acid were added to each tube, tubes were shaken manually for 10 seconds followed by centrifugation for 5 minutes at 20000g to separate phases. The lower phase was transferred to a fresh tube and tubes were dried in a speed-vac for 15 minutes at 45°C. Dried samples were redissolved by adding 1mL of spray buffer (8/5/1 isopropanol/methanol/H_2_O (all LC-MS grade) + 10mM ammonium acetate + 0.1% acetic acid (LC-MS grade)) to each sample and sonicating for 5 minutes in a bath sonicator. Mass spectra were recorded on a Thermo Q-Exactive Plus spectrometer equipped with a standard heated electrospray ionization (HESI) II ion source for shotgun lipidomics. Samples were sprayed at a flow rate of 10 µl min^−1^ in spray buffer. MS1 spectra (res. 280000) were recorded as segmented scans with windows of *m*/*z* 100 between *m*/*z* 240 and 1,200 followed by MS2 acquisition (res. 70000) from *m*/z 244.3364 to 1,199.9994 at *m*/*z* 1.0006 intervals. Raw files were converted into mzml files and analyzed using the LipidXplorer (version 1.2.8) (62). Internal standard intensities were used to calculate absolute amounts and all identified lipids were normalized to total lipid amount.

LCB levels measurements were carried out as contract work by the company Lipotype (Dresden, Germany). Data represent C14(dh)S plus C16(dh)S and C14(dh)S1P plus C16(dh)S1P of single animals. Because *cerS^P61^* are slimmer and smaller than control animals all values were normalized to the wet weight.

### Mechanonociception assays

Mechanonociception experiments were performed on staged 96 h AEL ± 3 h old 3rd instar larvae with a calibrated von-Frey-filament (50 mN). Larvae were carefully transferred with a wet brush to a 2% agar plate containing a 1 ml water film. Larvae were stimulated on mid-abdominal segments (A3-A5) twice within 2 s. Behavioral responses (non-nociceptive, bending, rolling/multiple rolling) to both stimuli were noted and only behavioral responses to the second stimulus were scored and plotted. Staging and experiments were done in a blinded and randomized fashion.

## Figure legends

**Suppl. Figure 1:**
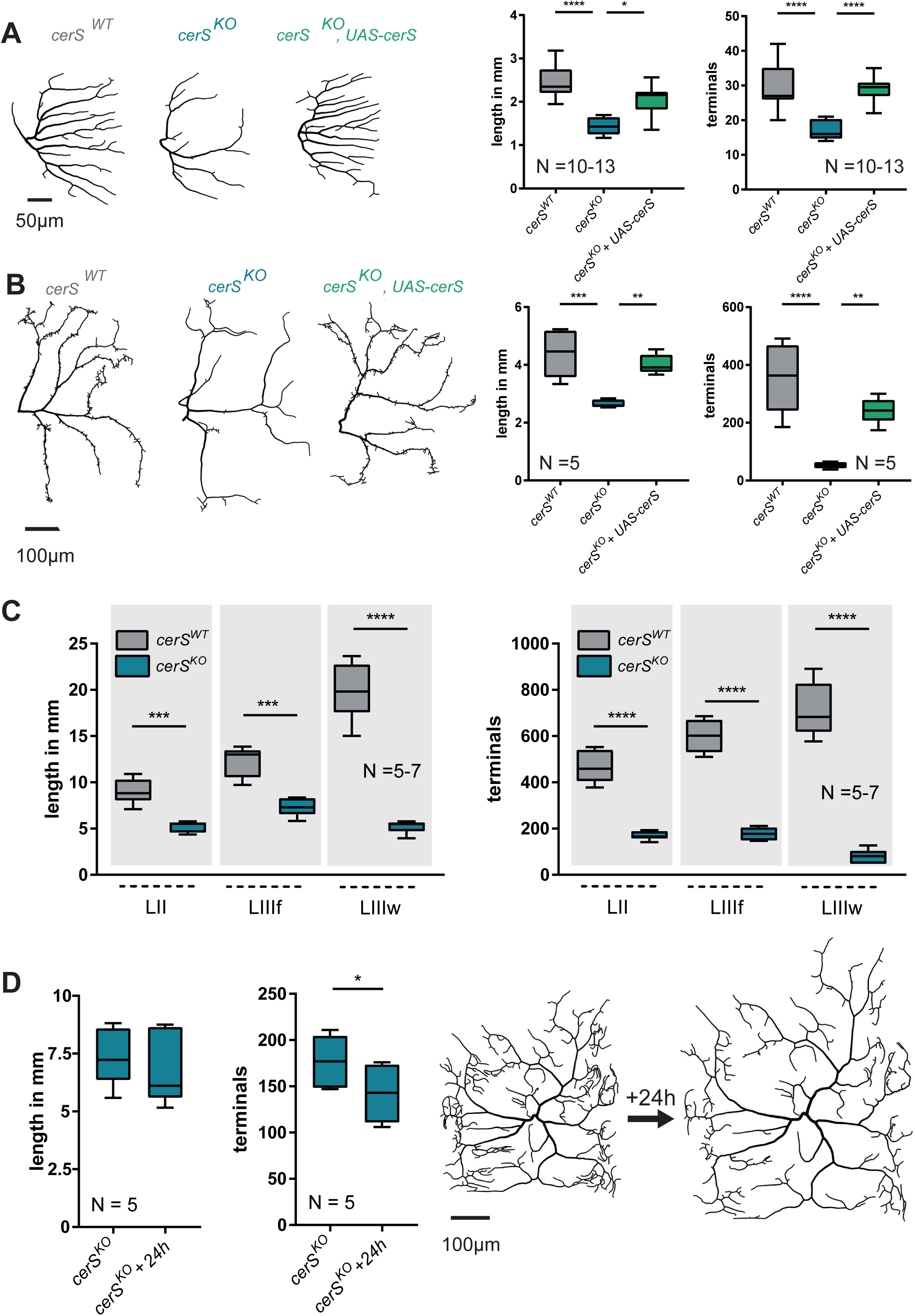
CerS activity is necessary for proper dendrite morphology in c1da and c3da neurons: Mutant c1da (ddaD, **A**) and c3da (ddaF, **B**) single cell clones expressing wild-type CerS (*cerS^WT^*) or no CerS (*cerS^KO^*) were generated using the MARCM technique. *cerS* expression was cell-autonomously rescued in *cerS^KO^* mutant neurons using a *UAS-cerS* expression construct (*cerS^KO^, UAS-cerS*). Representative tracings of the single-cell MARCM clones. Graphs show the quantified total dendrite length (left) and total number of dendritic terminal endpoints (right), respectively. Statistical test used: 1-Way-ANOVA followed by Dunnett’s multiple comparison test. **C**, Dendrites of mutant (*cerS^KO^*) and control (*CerS^WT^*) c4da neuronal MARCM clones were imaged at the second (LII), the feeding third (LIIIf), and the wandering third instar larval stage (LIIIw). Graphs show the quantified total dendritic length and the total number of terminal endpoints. Statistical test used: 2-Way-ANOVA followed by Sidak’s multiple comparison test. **D**, *cerS^KO^* c4da neuronal MARCM were imaged in early LIIIf stage larvae and 24h later. Images show representative dendritic trees and graphs the quantified total dendrite length and total number of dendritic terminal endpoints, respectively. Statistical test used in D: paired student’s *t*-test.

**Suppl. Figure 2:**
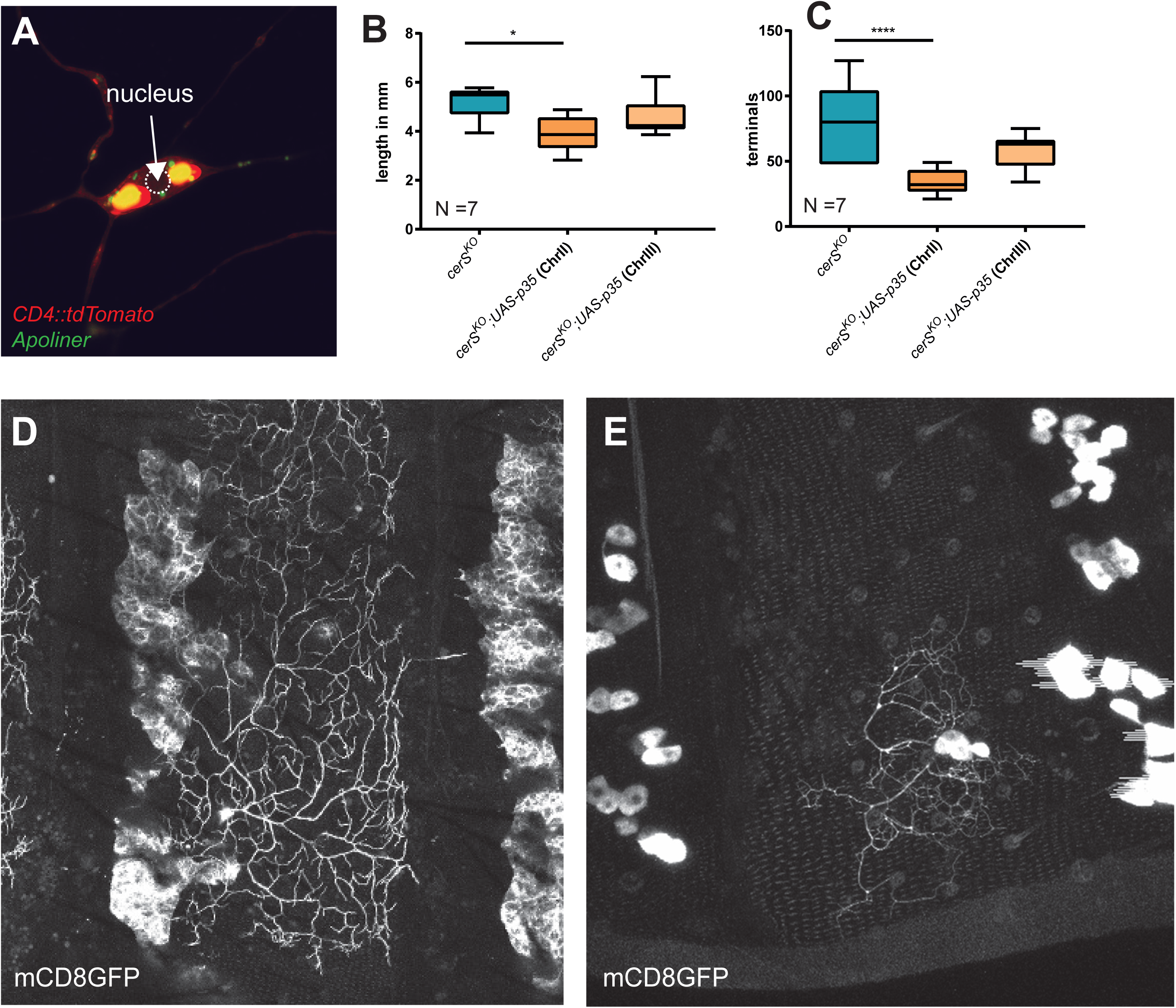
Morphological dendrite degeneration c4da neurons: **A,** *cerS^KO^* mutant MARCM clones were labeled by *UAS-CD4tdTomato* (red) and *UAS-Apoliner* expression is shown in green. Absence of nuclear GFP localization (arrow) indicates the lack of caspase activity. **B and C,** *cerS^KO^* mutant MARCM clones expressing caspase inhibitor p35 on the second or third chromosome do not show a rescue of total dendritic length (B) or number of total dendritic endpoints (C). **D and E,** dendrite morphology of wildtype (D) and *cerS^H215D^* mutant (E) c4da neuronal MARCM clones in late pupa (P14/P15). Statistical test used in B: paired student’s *t*-test and in C: 1-Way-ANOVA followed by Dunnett’s multiple comparison test.

**Suppl. Figure 3:**
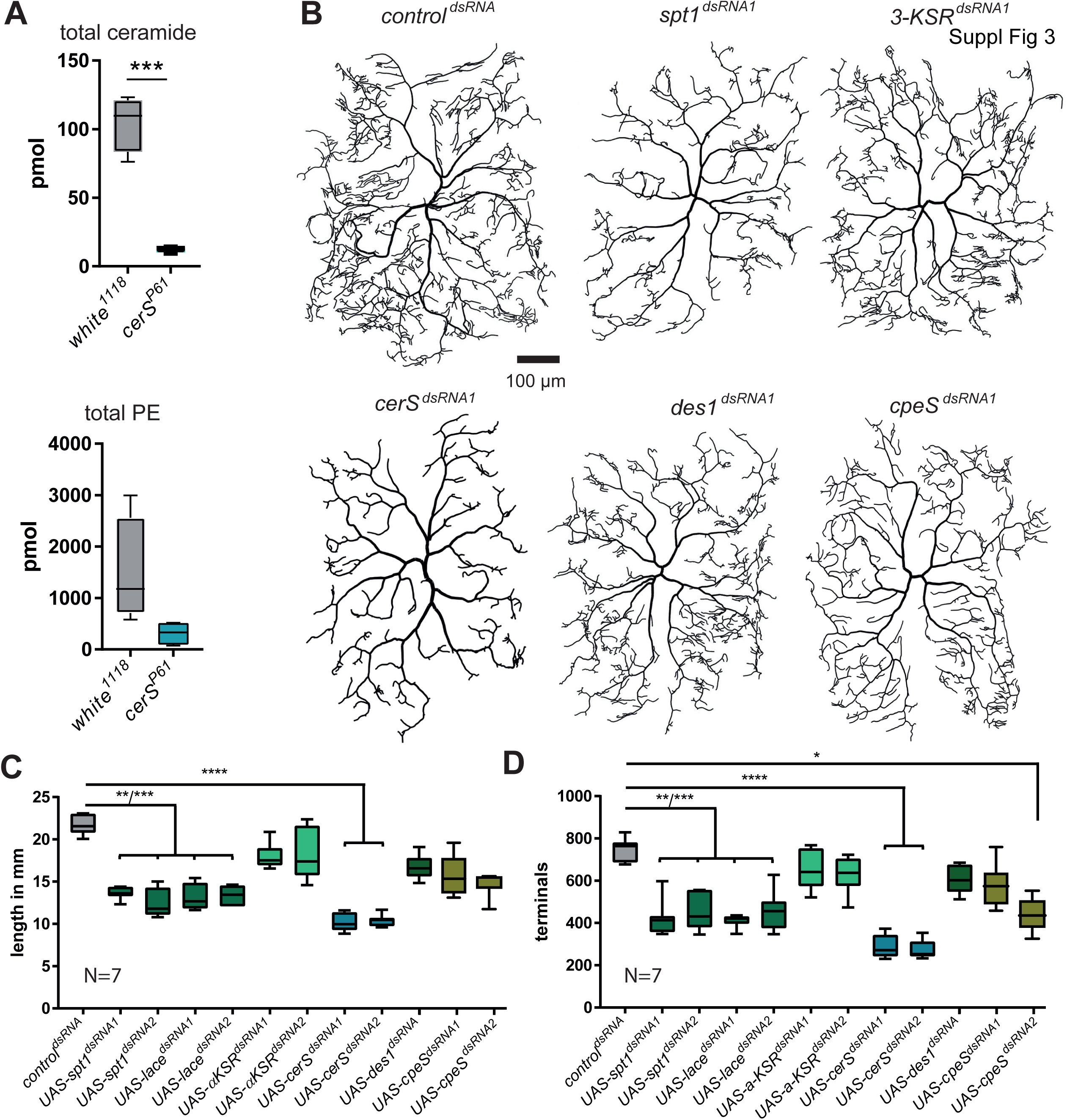
*cerS* knock-down strongly affects dendrite morphology: **A,** Total lipid levels and total PE levels in *white^1118^* and *cerS^P61^* mutant larvae. **B-D,** Knockdown of *spt1*, *lace, 3-KSR*, *cerS*, *des1* and *ceps* in c4da neurons was obtained using the c4da neuron specific ppk-Gal4 driver line and, if available, two independently generated UAS-dsRNA lines. C4da neurons were visualized by simultaneous co-expression of UAS-mCD8GFP. Representative dendrite tracings are shown in **B**. Quantified total dendrite length and total number of dendritic terminal endpoints are shown in **C** and **D**, respectively. Statistical test used in B: 1-Way-ANOVA followed by Dunnett’s multiple comparison test and C: Kruskal-Wallis with Dunńs multiple comparison test.

**Suppl. Figure 4:**
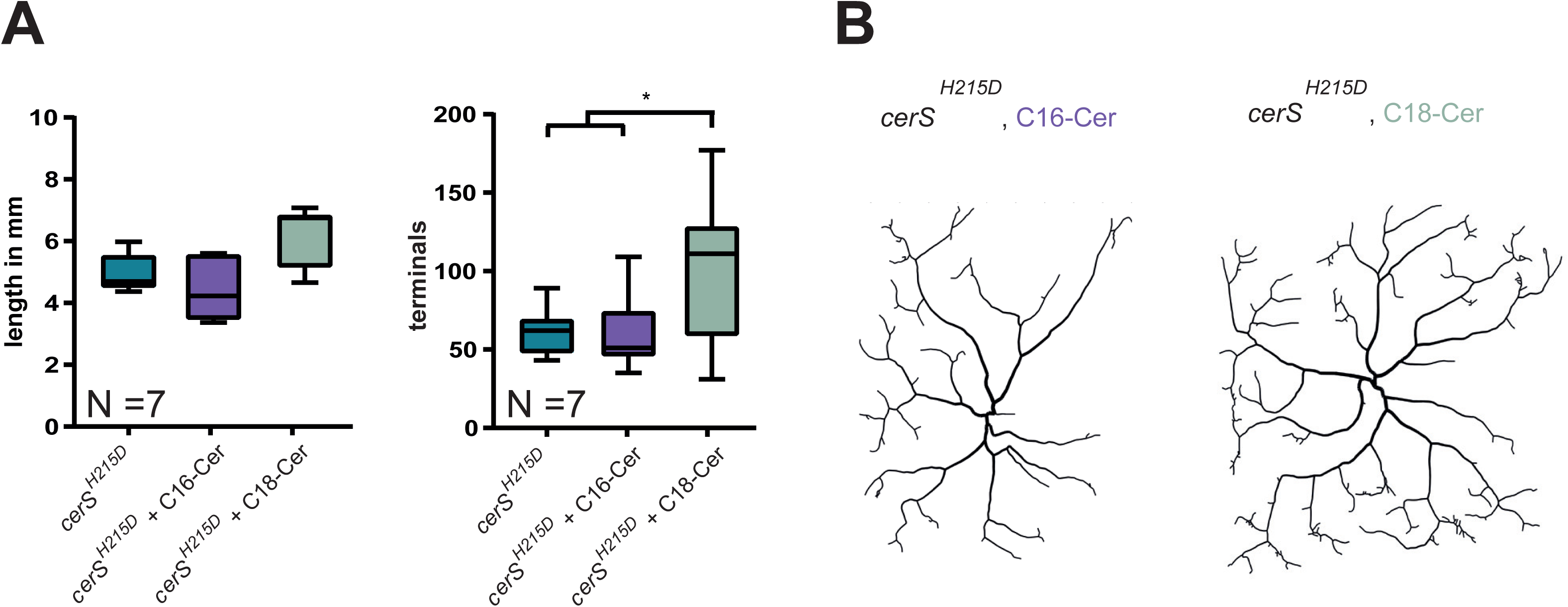
Dendrite defects are only mildly rescued by a C18-Cer enriched diet. **A/B**, larvae were raised on either C16- or C18-ceramide enriched food. Homozygous *cerS^H215D^* mutant c4da neurons were generated using the MARCM technique. Quantified total dendrite length and total number of dendritic terminal endpoints are shown in **A**. Representative neuronal tracings are shown in **B.** Statistical test used in A: 1-Way-ANOVA followed by Dunnett’s multiple comparison.

## Acknowledgements

We are grateful to Usha Acharya (University of Massachusetts Medical School, Worces, US) for sharing UAS-SK1 and UAS-SK2 flies and Corinne Antignac (Institute Imagine, Paris, France) for UAS-Sply flies. We thank Damian Demarest and Pia Bayer for contributing preliminary results and Mario Werner and Judith Reisen for help with cloning of UAS-CerS2, UAS-CerS-4, and UAS-CerS5 transgenes. We thank Karolina Doubkova for critical reading of the manuscript and Rita Kerpen, Regina Hube, Astrid Fleige, and Franka Eckhardt for Technical Assistance. This work was supported by institutional DZNE funding to GT and DFG ZI1690/2-1 to AZ.

